## Supplementary Materials for "Structural Adaptation of Fungal Cell Wall in Hypersaline Environment"

### Table of Contents


Supplementary Figure 1. Structural comparison of *A. sydowii* and *A. fumigatus*. 3

Supplementary Figure 2. Representative TEM images of *A. sydowii* 4

Supplementary Figure 3. Reproducibility of three batches of samples 5

Supplementary Figure 4. Changes in mobile carbohydrates in response to high salinity 6

Supplementary Figure 5. Intermolecular contacts between polysaccharides 7

Supplementary Figure 6. Water-edited experiments for examining polymer hydration 8

Supplementary Figure 7. NMR relaxation curves of polysaccharides 9

Supplementary Figure 8. Proteins and lipids mainly reside in the mobile phase 10

Supplementary Figure 9. Protein signals of rigid and mobile phases 11

Supplementary Figure 10. 2D $^{1}$H-$^{13}$C refocused INEPT spectra of phospholipids 12

Supplementary Figure 11. Membrane and lipid components in *A. sydowii* 13

Supplementary Table 1. Molar composition of rigid polysaccharides in *A. sydowii* cell wall 14

Supplementary Table 2. Intermolecular interactions identified by ssNMR 15

Supplementary Table 3. Water-edited intensities of polysaccharides cross peaks 16

Supplementary Table 4. $^{13}$C-$T_1$ and $^{1}$H-$T_{1\rho}$ time constants of polysaccharides 17

Supplementary Table 5. Water-edited intensities of amino acid residues 18

Supplementary Table 6. $^{13}$C and $^{1}$H chemical shifts of membrane lipids 19

Supplementary Table 7. Recipe of mineral-base liquid medium 20

Supplementary Table 8. Solid-state NMR experiments and parameters 21

Supplementary Table 9. $^{13}$C and $^{15}$N chemical shifts of *A. sydowii* polysaccharides and proteins 22

Supplementary References 23

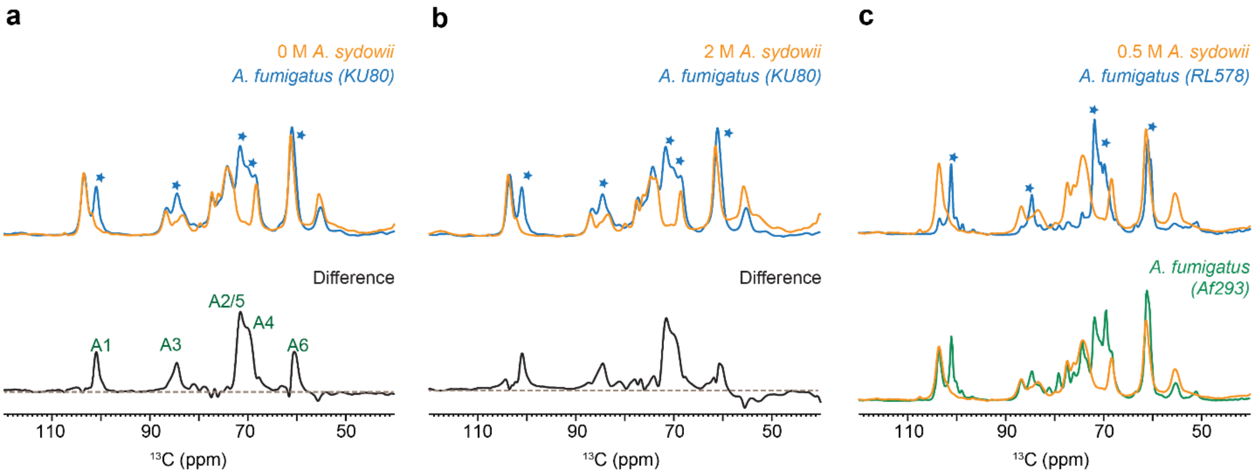


**Supplementary Figure 1. Structural comparison of *A. sydowii* and *A. fumigatus*.** Rigid polysaccharides shown by 1D $^{13}$C CP spectra of *A. fumigatus* KU80 (cyan) cultured at 0.1 M NaCl and *A. sydowii* (yellow) cultured at **a**, 0 M NaCl **b**, 2.0 M NaCl. Asterisks indicate the positions where the peak intensities were low in the *A. sydowii* sample. Subtraction of two parental spectra generates a difference spectrum showing α-1,3-glucan (A) signals, which are missing in *A. sydowii*. **c**, Overlay of 0.5 M NaCl *A. sydowii* with non-halophilic fungi *A. fumigatus* RL578 (top) and Af293 (bottom).

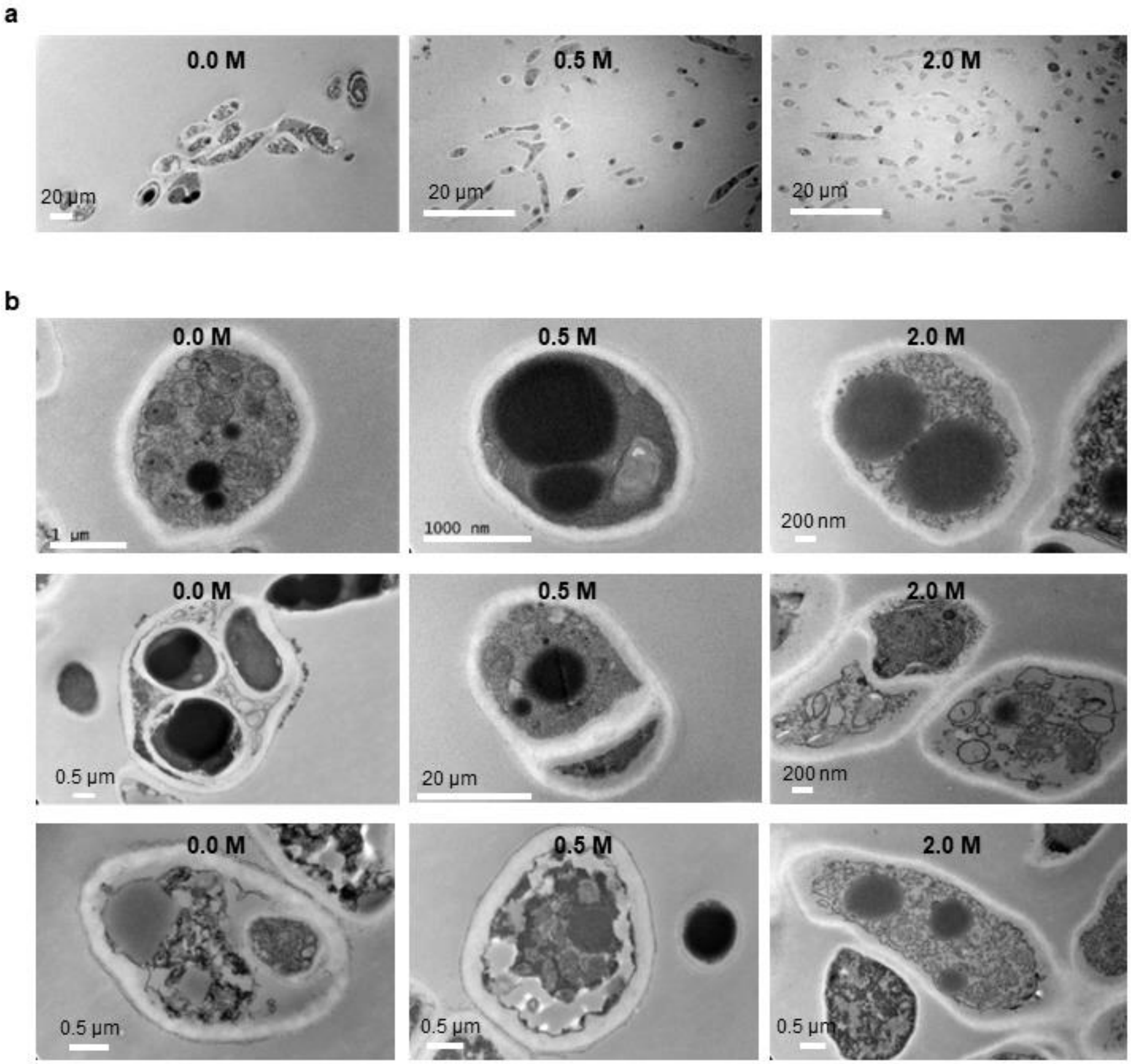


**Supplementary Figure 2. Representative TEM images of *Aspergillus sydowii*.** The images were taken from the perpendicular cross-sections of *A. sydowii* hyphae under **a,** low-magnification and **b,** high-magnification. The three columns of figures from the left to the right show the images of *A. sydowii* samples cultured at 0 M, 0.5 M, and 2 M NaCl conditions. White bars indicate the scale.

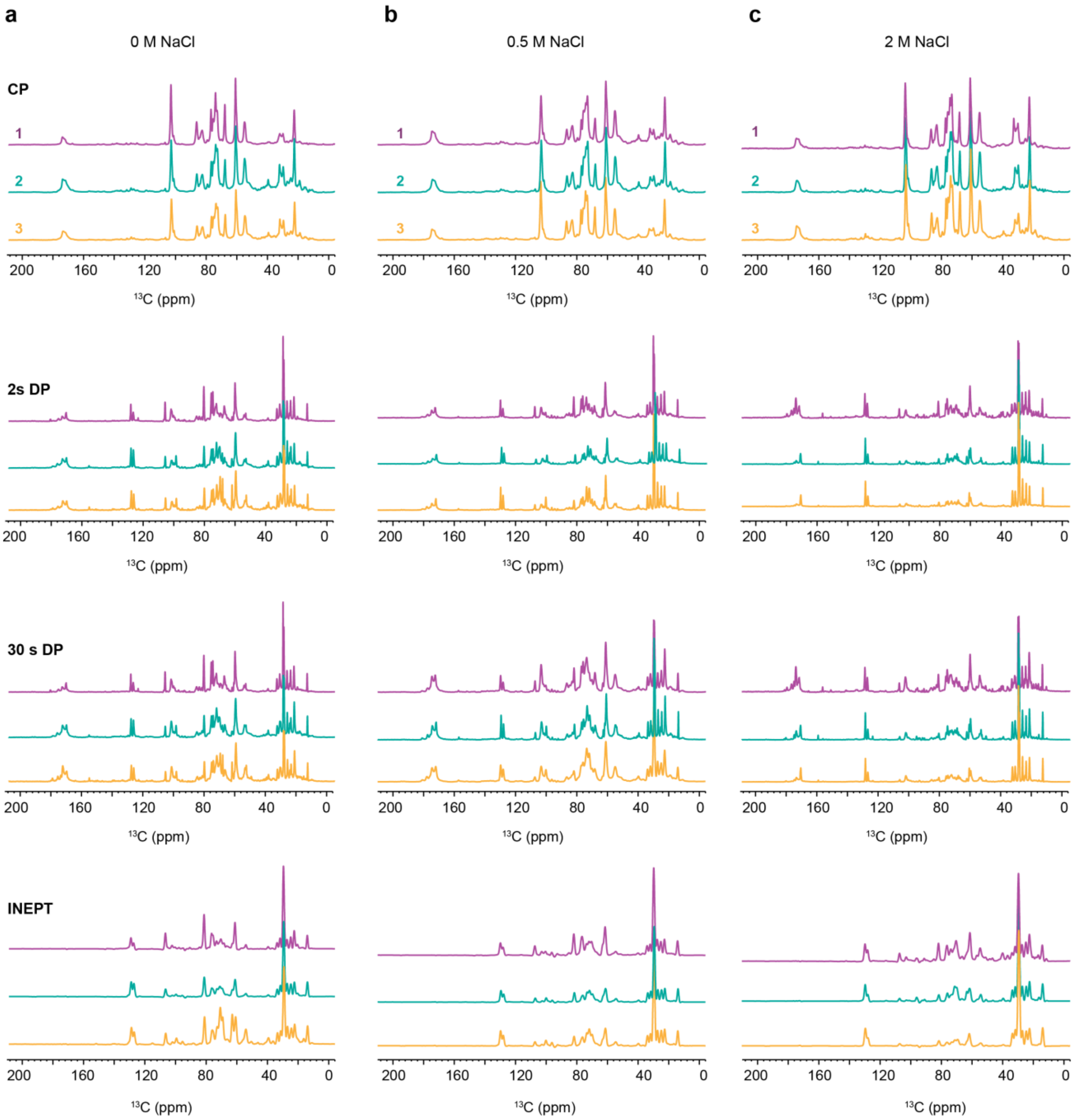


**Supplementary Figure 3. Reproducibility of three batches of *A. sydowii* samples.** 1D $^{13}$C spectra were collected on three batches of halophilic fungi cultured in **a**, 0 M, **b,** 0.5 M , and **c**, 2 M NaCl. 1D $^{13}$C CP spectra detecting the rigid molecules. 1D quantitative DP spectra were collected using a long recycle delay of 30 s for the unbiased detection of all molecules. 1D 2 s DP spectra detecting mobile molecules. 1D $^{13}$C refocused INEPT. Three batches of replicates labeled as 1 (purple), 2 (green), and 3 (yellow). Samples were measured in 800 MHz spectrometer at 13 kHz MAS at 298 K.

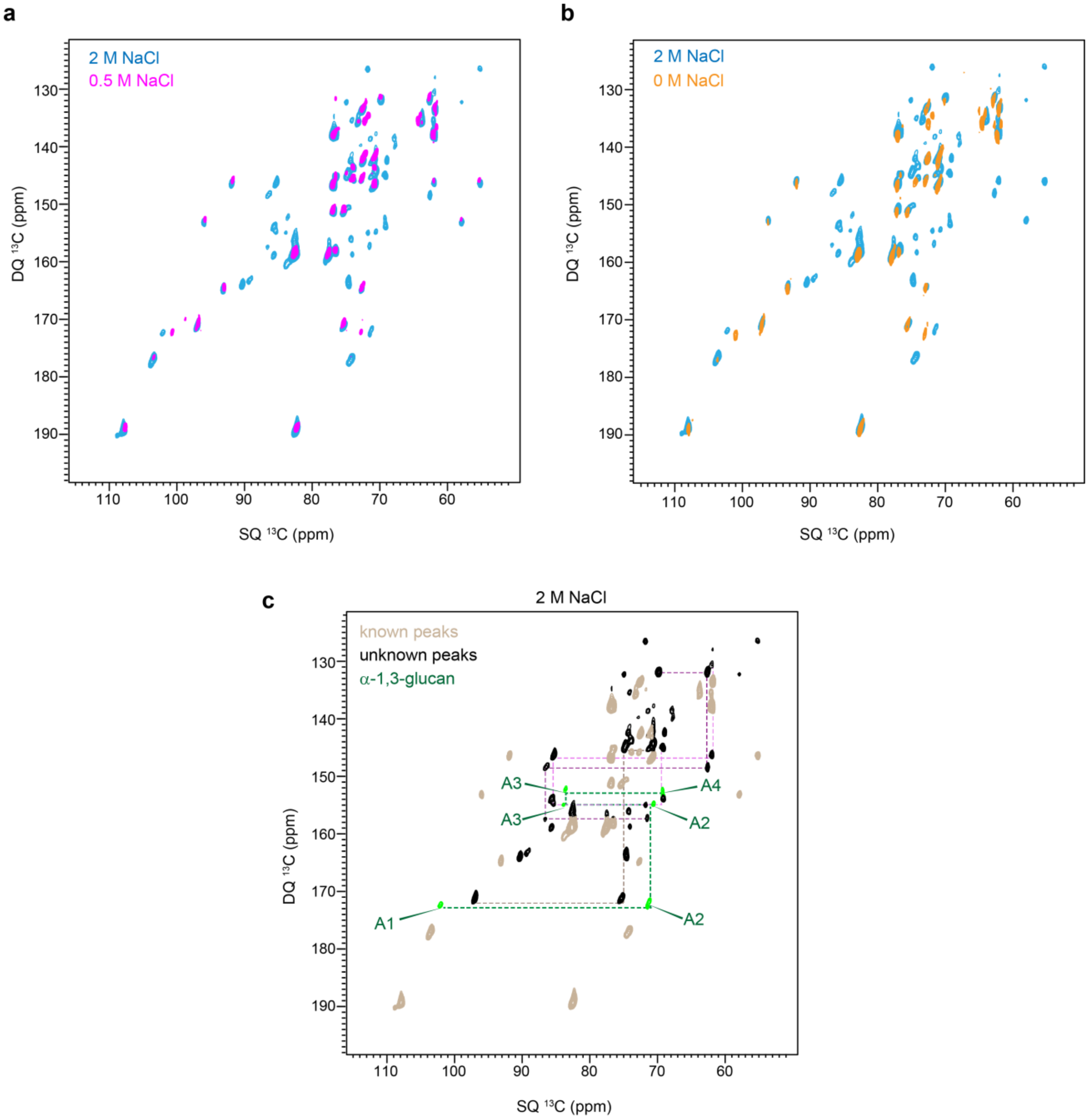


**Supplementary Figure 4. Changes in mobile carbohydrate in response to high salinity. a,** Overlay of 2D $^{13}C$ DP J-INADEQUATE spectra of *A. sydowii* samples cultured in 0.5 M (magenta) and 2 M (cyan) NaCl. **b,** Overlay of 2D $^{13}C$ DP J-INADEQUATE spectra of *A. sydowii* samples cultured in 0 M (orange) and 2 M (cyan) NaCl. **c,** 2D $^{13}C$ DP J-INADEQUATE spectra of 2 M NaCl showing the new peaks and α-1,3-glucan peaks. All spectra were measured in 850 MHz at 13 kHz MAS.

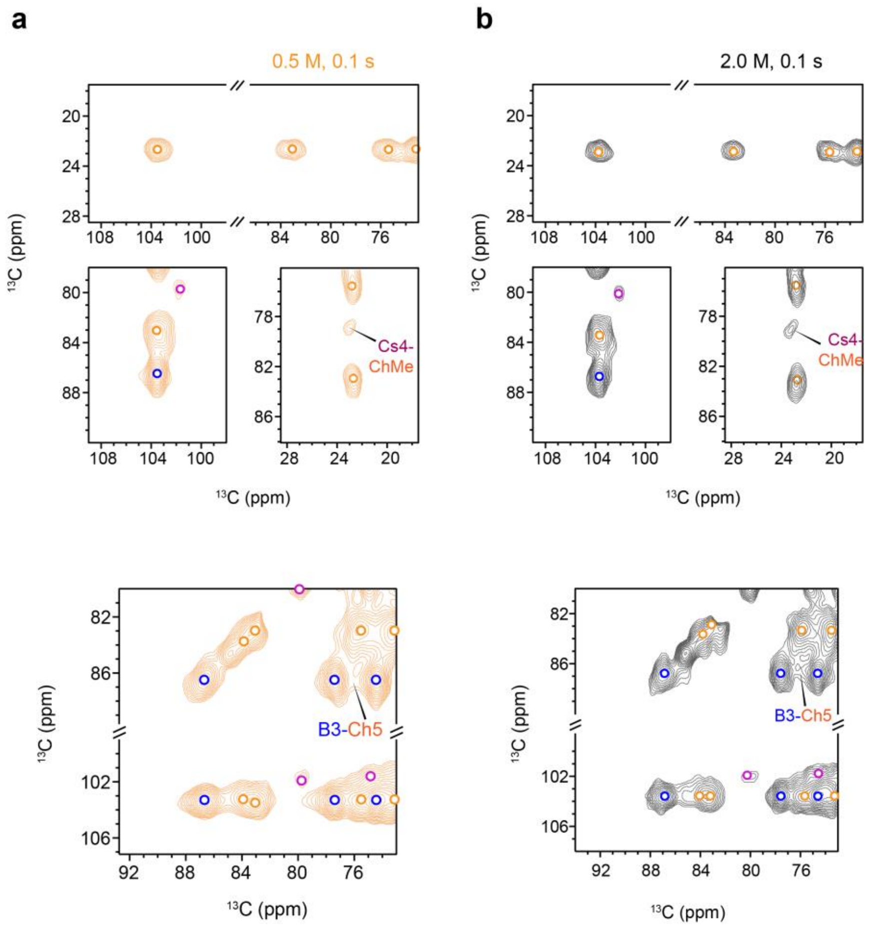


**Supplementary Figure 5. Intermolecular contacts between *A. sydowii* polysaccharides.** Selected regions were compared between **a,** 0.1 s DARR spectrum of sample cultured at 0.5 M NaCl, **b,** 0.1 s DARR spectrum of sample cultured at 2.0 M NaCl. and Open circles indicate the positions of intramolecular signals within chitin (orange), β-glucan (blue), and chitosan (purple). Intermolecular interactions are labeled. For example, Cs4-ChMe represents the cross peak between chitin methyl carbon and chitosan carbon 4. All spectra were measured on an 850 MHz NMR spectrometer.

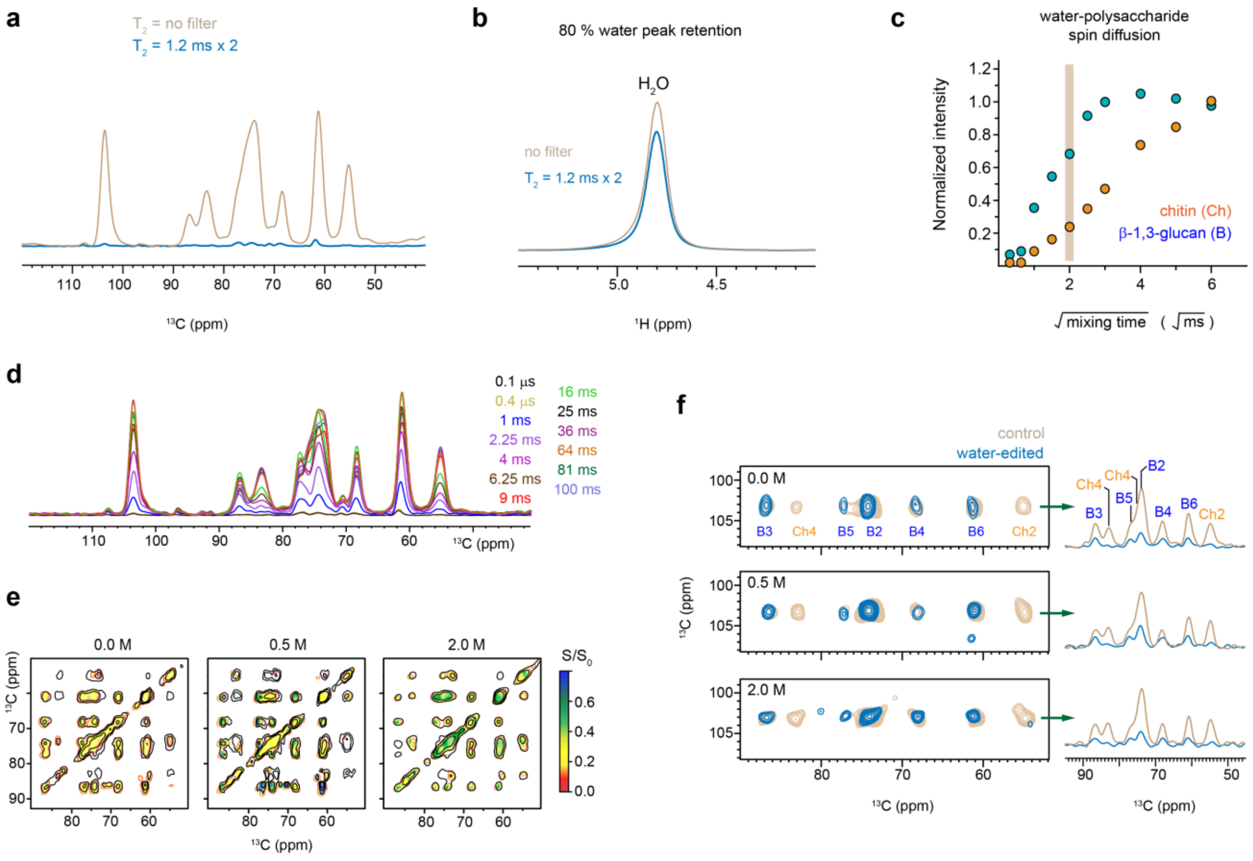


**Supplementary Figure 6. Water-edited experiments for examining polymer hydration. a**, Overlay of $^{13}C$ CP spectra of *A. sydowii* 0.5 M sample (in almond) with a $^{1}H$ $T_2$ filtered spectrum ($T_2$ = 1.2 ms x 2, blue). No spin diffusion was applied. 96% of carbohydrate signals were removed by the filter. **b**, 80% of water magnetization was retained after the $^{1}H$-$T_2$ filter. **c**, Representative water-to-polysaccharide $^{1}H$ spin diffusion build-up curves. β-1,3-glucan has a faster buildup curve than chitin, revealing the hydrophilic nature of β-1,3 glucan. The curves are obtained from peak intensities of water-edited $^{13}C$ spectra. **d**, 1D water-edited $^{13}C$ spectra with different $^{1}H$ mixing times. **e**, Heatmap of relative water-edited intensity ratios ($S/S_0$) of *A. sydowii* cell wall polysaccharides presented as a 2D $^{13}C$-$^{13}C$ correlation spectra. **f**, Overlay of 2D water-edited spectra (blue) and control spectra (almond). The β-1,3-glucan peaks were selectively retained due to the water-association of this polysaccharide. 1D $^{13}C$ cross sections were extracted for comparison. All the spectra were measured on a 400 MHz spectrometer at 10 kHz MAS.

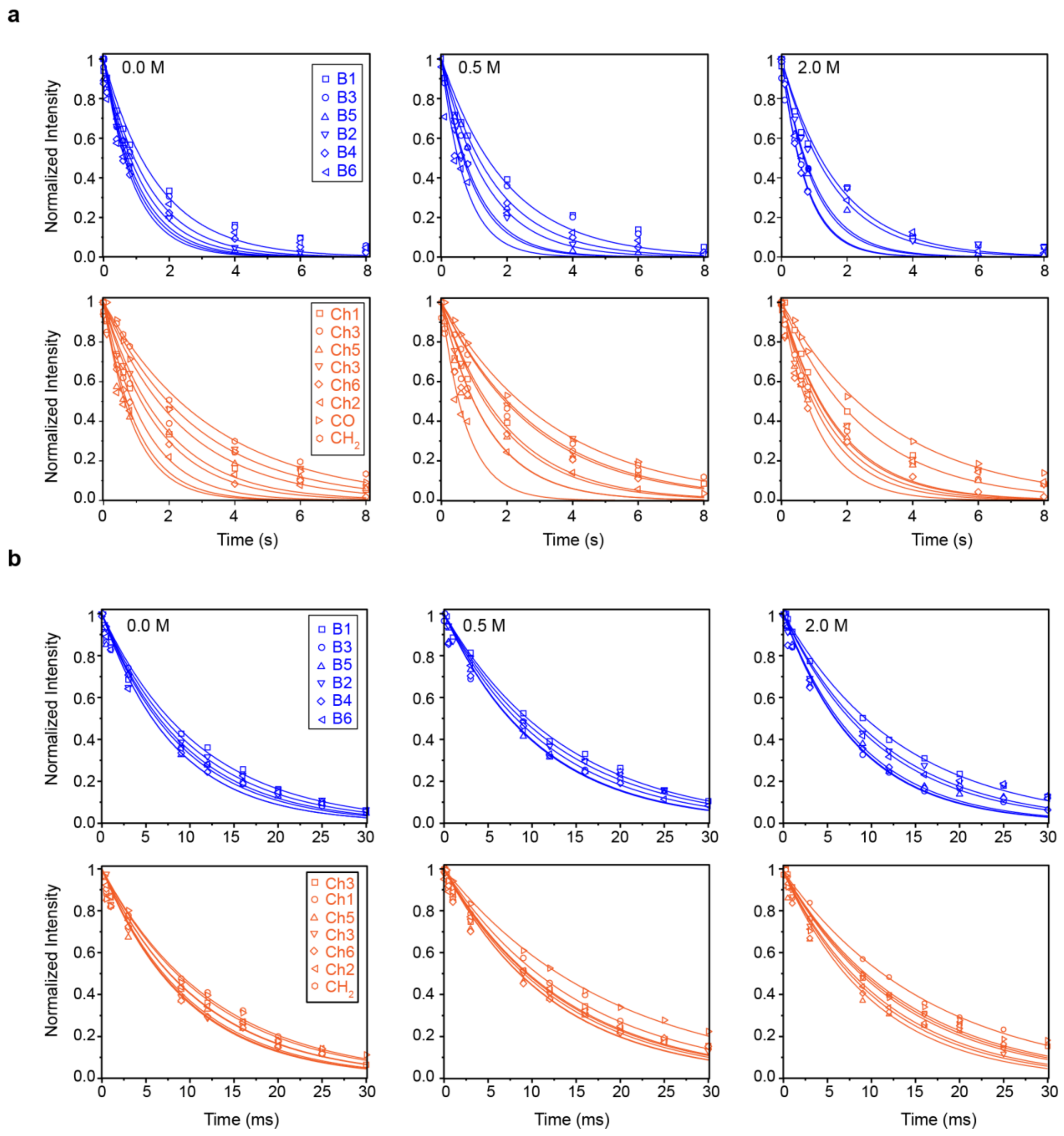


**Supplementary Figure 7. NMR relaxation curves of polysaccharides. a,** $^{13}C$-$T_1$ and **b,** $^{1}H$-$T_{1\rho}$ relaxation curves of polysaccharides in intact *A. sydowii* cell wall across 0 M, 0.5 M and 2 M NaCl. The data are collected on 400 MHz (9.4 Tesla) spectrometer at 10 kHz MAS and the best fit is achieved using single exponential equation. Blue curves are β-1,3-glucan and orange curves are chitin. Symbols are used for assigning different carbons in the polysaccharides.

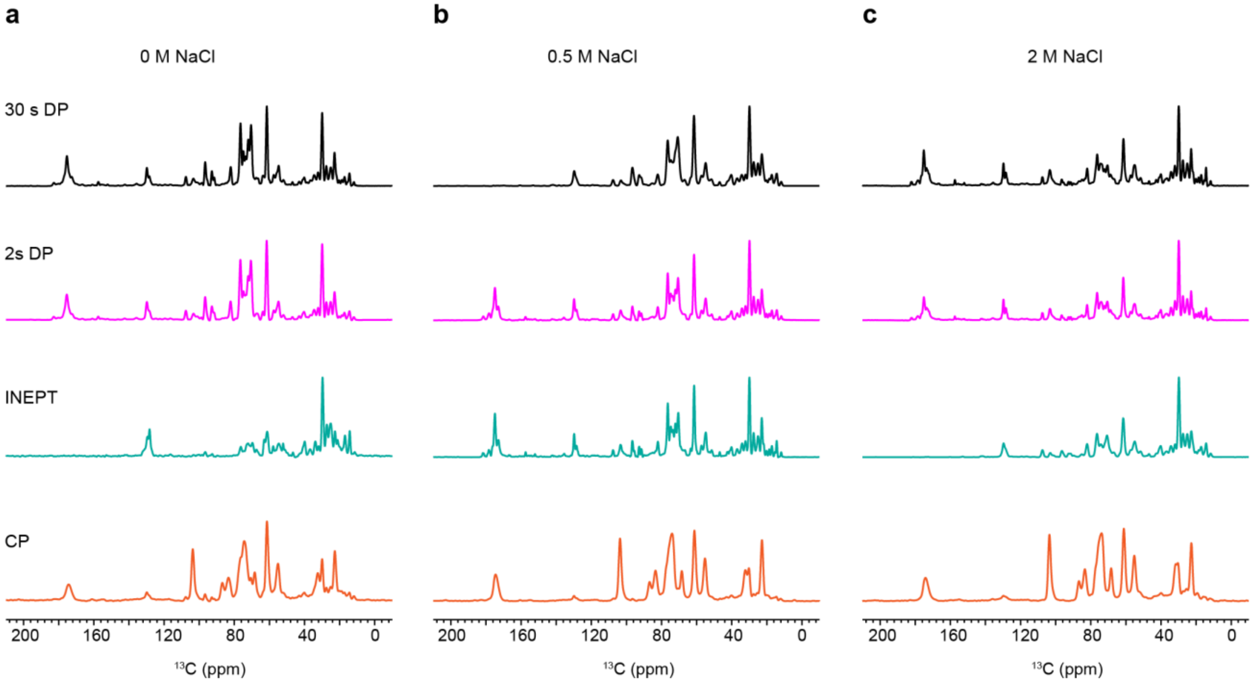


**Supplementary Figure 8.** ***A. sydowii*** **proteins and lipids mainly reside in the mobile phase.** An array of 1D $^{13}$C spectra that detect different components with distinct dynamics are compared for *A. sydowii* samples cultured with **a,** 0 M, **b,** 0.5 M, and **c,** 2 M NaCl. These spectra include the 1D $^{13}$C DP spectra measured with long recycle delays of 30 s (quantitative detection) and short recycle delays of 2 s (preferential detection of mobile molecules), 1D INEPT spectra (very mobile components) and 1D CP spectra (rigid molecules). All the spectra were measured on 400 MHz NMR under 10 kHz MAS.

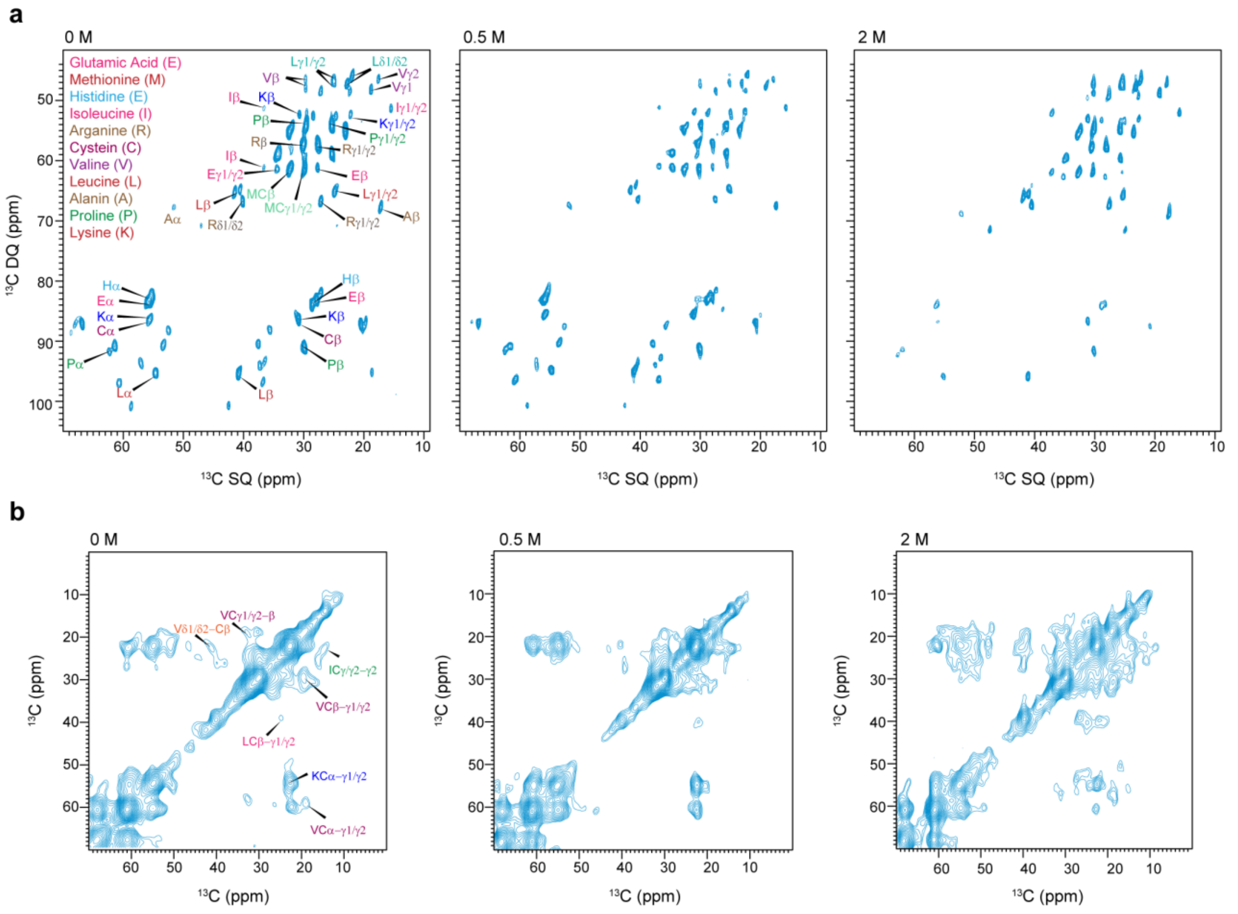

**Supplementary Figure 9. Protein signals of mobile and rigid phases. a,** 2D $^{13}C$ DP refocused J-INADEQUATE spectra showing signals of mobile proteins. **b,** 2D $^{13}C$-$^{13}C$ CP-based 100 ms DARR spectra showing the rigid proteins of 0 M, 0.5 M, and 2 M samples. All the refocused INADEQUATE spectra were measured on a 850 MHz spectrometer at 13 kHz MAS and all the DARR spectra were measured on a 400 MHz spectrometer at 10 kHz MAS.

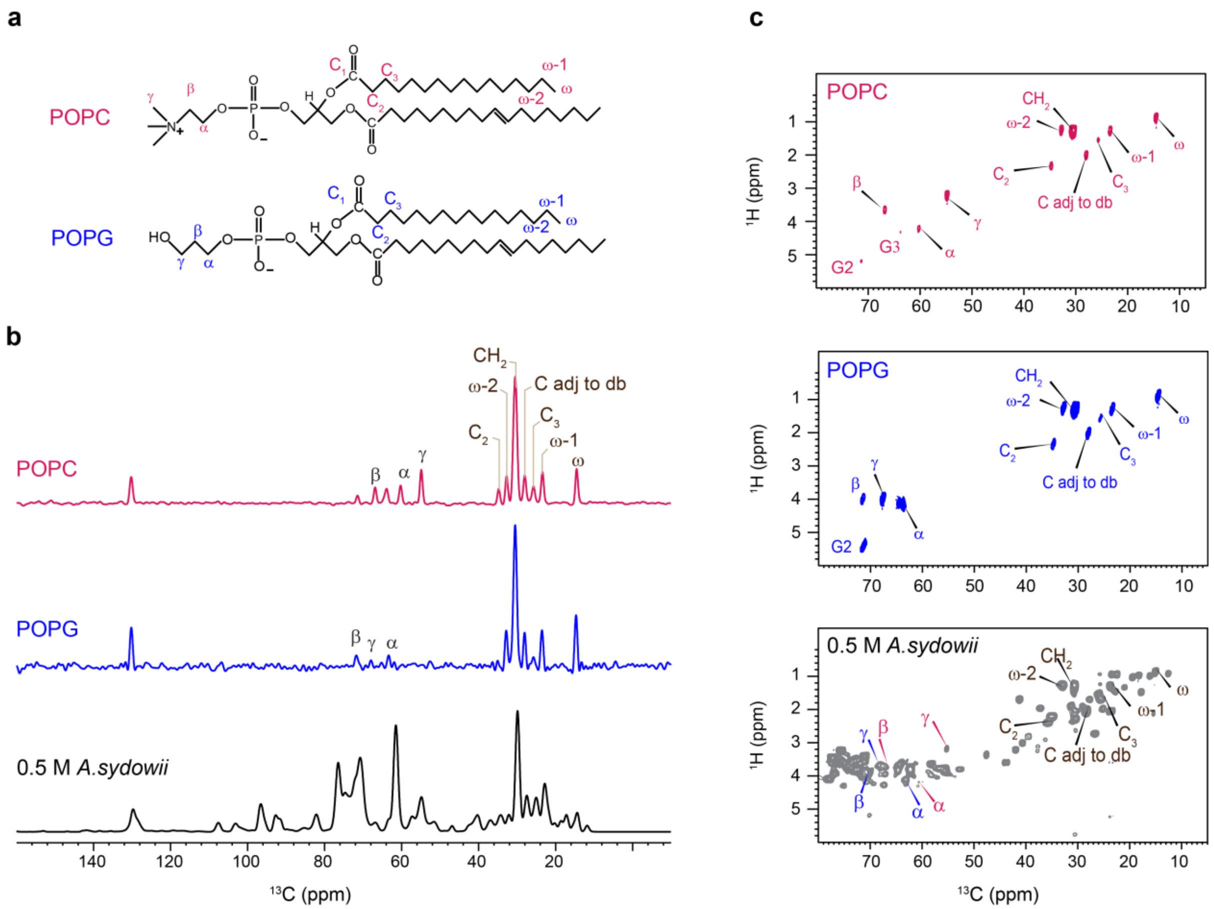


**Supplementary Figure 10. 2D $^{1}$H-$^{13}$C refocused INEPT spectra of phospholipids. a,** Chemical structure of model phospholipids POPC and POPG with carbons labeled. **b,** 1D $^{13}$C INEPT spectra and **c,** 2D $^{1}$H-$^{13}$C spectra of POPC, POPG and *A. sydowii* with 0.5 M NaCl. The spectra were measured in 400 MHz spectrometer at 10 kHz MAS.

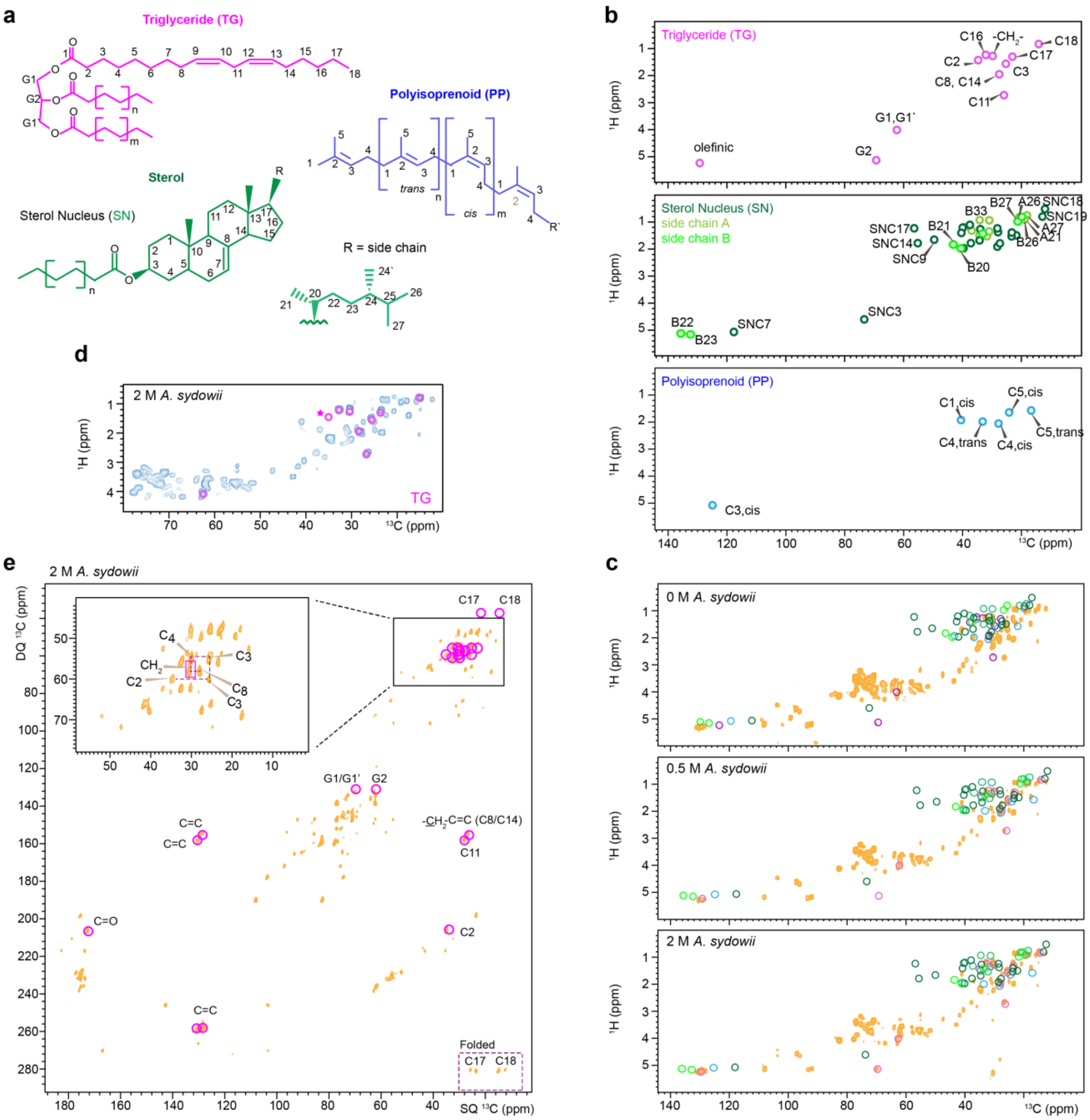

**Supplementary Figure 11. Membrane and lipid components in *A. sydowii*. a,** Representative structures of three lipid components: triglycerides (TG), Sterol (S), and polyisoprenoid (PP). **b,** Simulated 2D $^{1}$H-$^{13}$C spectra using literature reported chemical shifts of TG, and S, and PP documented in **Supplementary Table 6**. **c,** Overlay of 2D $^{1}$H-$^{13}$C refocused INEPT spectra of 0 M, 0.5 M and 2.0 M samples with stimulated spectra. The spectra of samples with 0 M and 0.5 M NaCl do not contain signals from triglycerides and sterols. **d,** Most TG signals are present in 2D $^{1}$H-$^{13}$C refocused INEPT spectrum of *A. sydowii* with 2.0 M NaCl except for the signals of C2 (highlighted by asterisk). **e,** Overlay of simulated TG signals and experimentally measured DP refocused *J*-INADEQUATE spectrum of 2 M *A. sydowii*. All expected signals are present, though the signals are heavily overlapped with other lipids, amino acids, as well as the C5-C6 of some carbohydrates. Note that the signals of the C17-C18 spin pair is folded to the bottom right corner of the spectrum due to limited window width of the spectrum.

**Supplementary Table 1. Molar composition of rigid polysaccharides in *A. sydowii* cell wall.** The numbers were calculated using the integrals of well-resolved cross peaks of β-1,3 glucan and chitin in 2D $^{13}C$-$^{13}C$ DARR spectra. The results were already normalized by the number of scans. Error bars are standard errors of cross peak intensities. Not detected (-).

| Rigid molecules | | | | |
|---|---|---|---|---|
| Polysaccharide | | 0 M NaCl | 0.5 M NaCl | 2 M NaCl |
| β-1,3-glucan | | 55 ± 8%[a] | 51 ± 9%[b] | 40 ± 7%[c] |
| Chitin | | 33 ± 7 %[d] | 39 ± 8%[e] | 50 ± 13%[f] |
| Chitosan | | 12 ± 3% | 10 ± 2% | 10 ± 3% |
| Mobile molecules | | | | |
| GAG | GalN | 4 ± 1% | 6 ± 2% | 13 ± 3% |
| | GalNAc | 4 ± 1% | 5 ± 1% | 9 ± 2% |
| | Gal*p* | 20 ± 6% | 24 ± 3% | 14 ± 7% |
| GM | | 62 ± 13% | 58 ± 15% | 43 ± 17% |
| β-1,3-glucan | | 11 ± 3% | 7 ± 2% | 14 ± 5% |
| α-1,3-glucan | | - | - | 6 ± 1% |

[a] Percentage taken from the average of B1-3, B1-5, B1-2, B1-4, B5-6, B3-4 cross peak integrations.
[b] The average of B1-3, B1-5, B1-2, B1-4, B3-5, B3-4 cross peak integrations.
[c] The average of B1-3, B1-5, B1-4, B3-2, B3-5, B3-4, B3-6, B5-4, B5-2 cross peak integrations.
[d] The average of Ch1-2, Ch1-6, Ch4-2, Ch5-2, Ch3-6, cross peak integrations.
[e] The average of Ch1-4, Ch1-6, Ch1-2, Ch4-6, Ch4-2, Ch5-2, cross peak integrations.
[f] The average of Ch1-4, Ch1-6, Ch1-3, Ch1-2, Ch4-2, Ch5-4, Ch5-2 cross peak integrations.

**Supplementary Table 2. Intermolecular interactions identified by ssNMR.** The table documented the cross peaks between different polysaccharides in *A. sydowii* samples cultured at 0.5 M and 2 M salinity conditions. The chemical shifts for the two dimensions of the spectra ($\omega_1$ and $\omega_2$), the assignment of the cross peak, the type of spectra, and the sample condition are summarized.

| Cross peak | $\omega_1$, $\omega_2$ (ppm, ppm) | 0.5 M 0.1 s PDSD | 0.5M 1.5 s PDSD | $\omega_1$, $\omega_2$ (ppm, ppm) | 2M 0.1 s PDSD | 2M 1.5 s PDSD |
|---|---|---|---|---|---|---|
| Ch4-Ch4’ | 82.3, 84.1 | | x | 82.3, 84.1 | | x |
| Ch4’-Ch4 | 84.1, 82.3 | | x | 84.1, 82.3 | | x |
| ChMe-B5 | 23.1, 77.1 | | x | 23.1, 77.1 | | x |
| B3-Ch5 | 86.1, 76.1 | x | x | 86.1, 76.1 | x | x |
| B5-Ch5 | 68.2, 75.5 | x | x | 68.5, 75.8 | x | x |
| ChMe-Cs4 | 22.9, 79.4 | | x | 22.7, 79.9 | | x |
| ChMe-Cs1 | 22.9, 101.5 | | x | 22.7, 102.0 | | x |
| Ch4-Cs1 | 83.5, 101.5 | | x | | | |
| Ch1/B1-Cs4 | 103.2, 79.5 | | x | | | |
| Cs4-ChMe | 79.2, 22.9 | x | x | 79.2, 22.8 | x | x |
| Cs4-ChCO | 79.7, 173.6 | | | 79.7, 173.8 | | x |
| Cs1-Ch4 | 102.1, 83.1 | | x | | | |
| B1-Cs1 | | | | 86.8, 102.1 | | x |
| B3-Cs4 | 86.5, 79.2 | | x | | | |
| B5-Cs4 | 68.2, 79.2 | | x | | | |
| Cs1-B5 | 101.5, 77.2 | | x | | | |

**Supplementary Table 3. Water-edited intensities of polysaccharides cross peaks.** The intensity ratios were obtained by comparing the peak intensity in water-edited and control spectra, with normalization by the number of scans. Error bars are standard deviations propagated from NMR signal-to-noise ratios.

| | 0.5 M NaCl | | 0 M NaCl | | 2 M NaCl | |
|---|---|---|---|---|---|---|
| | **Cross peak** | **Intensity** | **Cross peak** | **Intensity** | **Cross peak** | **Intensity** |
| β-1,3-glucan | B1-3 | 0.5±0.1 | B1-3 | 0.42±0.09 | B1-3 | 0.4±0.1 |
| | B1-5 | 0.5±0.1 | B1-5 | 0.37±0.09 | B1-5 | 0.5±0.1 |
| | B1-2 | 0.41±0.04 | B1-2 | 0.26±0.03 | B1-2 | 0.31±0.04 |
| | B1-4 | 0.5±0.1 | B1-4 | 0.29±0.07 | B1-4 | 0.5±0.1 |
| | B1-6 | 0.39±0.07 | B1-6 | 0.28±0.06 | B1-6 | 0.34±0.09 |
| | B3-1 | 0.7±0.1 | B3-1 | 0.43±0.07 | B3-1 | 0.5±0.1 |
| | B3-2 | 0.61±0.09 | B3-5 | 0.34±0.06 | B3-5 | 0.4±0.1 |
| | B3-4 | 0.5±0.1 | B3-2 | 0.44±0.06 | B3-2 | 0.3±0.1 |
| | B3-6 | 0.80±0.07 | B3-4 | 0.38±0.07 | B3-4 | 0.4±0.1 |
| | B5-1 | 0.45±0.08 | B3-6 | 0.59±0.06 | B3-6 | 0.5±0.2 |
| | B5-3 | 0.7±0.2 | B5-1 | 0.35±0.07 | B5-1 | 0.4±0.1 |
| | B5-2 | 0.40±0.07 | B5-3 | 0.4±0.1 | B5-3 | 0.5±0.2 |
| | B5-4 | 0.44±0.08 | B5-2 | 0.25±0.07 | B5-2 | 0.21±0.07 |
| | B5-6 | 0.39±0.05 | B5-4 | 0.28±0.06 | B5-4 | 0.36±0.07 |
| | B2-1 | 0.39±0.04 | B5-6 | 0.30±0.05 | B5-6 | 0.37±0.06 |
| | B2-3 | 0.6±0.1 | B2-1 | 0.29±0.03 | B2-1 | 0.25±0.04 |
| | B2-5 | 0.7±0.1 | B2-3 | 0.39±0.09 | B2-3 | 0.6±0.2 |
| | B2-4 | 0.6±0.1 | B2-5 | 0.53±0.09 | B2-5 | 0.5±0.1 |
| | B2-6 | 0.31±0.04 | B2-4 | 0.36±0.07 | B2-4 | 0.38±0.07 |
| | B4-1 | 0.6±0.1 | B2-6 | 0.25±0.04 | B2-6 | 0.26±0.05 |
| | B4-3 | 0.6±0.1 | B4-1 | 0.36±0.07 | B4-1 | 0.5±0.1 |
| | B4-5 | 0.6±0.1 | B4-3 | 0.37±0.09 | B4-3 | 0.3±0.1 |
| | B4-2 | 0.43±0.09 | B4-5 | 0.38±0.07 | B4-5 | 0.4±0.1 |
| | B4-6 | 0.5±0.1 | B4-2 | 0.42±0.08 | B4-2 | 0.28±0.08 |
| | | | B4-6 | 0.26±0.07 | B4-6 | 0.33±0.09 |
| Chitin | Ch1-3 | 0.13±0.02 | Ch1-4 | 0.21±0.02 | Ch1-4 | 0.14±0.09 |
| | Ch1-6 | 0.38±0.07 | Ch1-5 | 0.38±0.05 | Ch1-3 | 0.23±0.04 |
| | Ch4-1 | 0.12±0.09 | Ch1-2 | 0.12±0.06 | Ch4-1 | 0.2±0.1 |
| | Ch4-4 | 0.23±0.08 | Ch4-4 | 0.22±0.08 | Ch5-1 | 0.27±0.08 |
| | Ch5-1 | 0.35±0.06 | Ch5-1 | 0.42±0.06 | Ch5-3 | 0.06±0.04 |
| | Ch5-4 | 0.15±0.8 | Ch5-4 | 0.17±0.09 | Ch5-6 | 0.19±0.5 |
| | Ch5-3 | 0.2±0.03 | Ch5-3 | 0.21±0.05 | Ch5-2 | 0.2±0.1 |
| | Ch5-6 | 0.27±0.05 | Ch5-6 | 0.31±0.04 | Ch3-1 | 0.26±0.07 |
| | Ch3-1 | 0.22±0.05 | Ch5-2 | 0.09±0.06 | Ch3-4 | 0.1±0.1 |
| | Ch3-5 | 0.12±0.01 | Ch3-1 | 0.31±0.05 | Ch3-6 | 0.29±0.09 |
| | Ch3-6 | 0.21±0.07 | Ch3-5 | 0.29±0.02 | Ch3-2 | 0.14±0.08 |
| | Ch3-2 | 0.13±0.07 | Ch3-6 | 0.22±0.06 | Ch2-1 | 0.2±0.1 |
| | Ch2-1 | 0.22±0.07 | Ch3-2 | 0.15±0.05 | Ch2-6 | 0.2±0.1 |
| | Ch2-4 | 0.2±0.1 | Ch2-1 | 0.17±0.09 | Ch2-2 | 0.1±0.04 |
| | Ch2-5 | 0.12±0.07 | Ch2-3 | 0.08±0.07 | | |
| | Ch2-3 | 0.13±0.06 | | | | |
| | Ch2-6 | 0.2±0.1 | | | | |

**Supplementary Table 4. $^{13}$C-$T_1$ and $^{1}$H-$T_{1\rho}$ relaxation time constants of polysaccharides in *A. sydowii*.** A single exponential equation was used to fit the $T_1$ data $I(t) = e^{-t/T_I}$. A single exponential equation was used to fit the $T_{1\rho}$ data: $I(t) = e^{-t/T_{I\rho}}$. Error bars are standard deviations of the fit parameters.

| Sample Type | Cross peaks | $T_1$ (s) | Cross peaks | $T_{1\rho}$ (ms) |
|---|---|---|---|---|
| 0.5 M NaCl | B3 | 1.5±0.1 | B1 | 13.8±0.8 |
| | B5 | 0.8±0.2 | B3 | 11±1 |
| | B2 | 1.6±0.3 | B5 | 11±1 |
| | B4 | 0.7±0.3 | B2 | 12.7±0.9 |
| | B6 | 0.5±0.1 | B4 | 11±1 |
| | Ch1 | 2.2±0.2 | B6 | 12±1 |
| | Ch5 | 1.0±0.2 | Ch1 | 13.8±0.8 |
| | Ch3 | 2.2±0.4 | Ch4 | 15.3±0.8 |
| | Ch6 | 0.8±0.1 | Ch5 | 12.4±0.9 |
| | Ch2 | 2.0±0.4 | Ch3 | 13.1±0.8 |
| | | | Ch6 | 12±1 |
| | | | Ch2 | 13.5±0.8 |
| 0 M NaCl | B1 | 1.7 ±0.2 | B1 | 11±0.6 |
| | B3 | 1.28±0.07 | B3 | 8.9±0.9 |
| | B5 | 0.9±0.1 | B5 | 8.3±0.8 |
| | B2 | 1.4±0.2 | B2 | 10.1±0.6 |
| | B4 | 0.88±0.05 | B4 | 8.3±0.8 |
| | B6 | 0.9±0.1 | B6 | 9.4±0.6 |
| | Ch1 | 1.7±0.2 | Ch1 | 11.0±0.6 |
| | Ch4 | 2.9±0.3 | Ch4 | 12.6±0.9 |
| | Ch5 | 1.1±0.2 | Ch5 | 9.8±0.9 |
| | Ch3 | 1.3±0.2 | Ch3 | 10.1±0.7 |
| | Ch2 | 1.5±0.3 | Ch6 | 9.7±0.6 |
| | | | Ch2 | 11.1±0.6 |
| 2.0 M NaCl | B1 | 1.7±0.2 | B1 | 13.5±0.6 |
| | B3 | 0.77±0.07 | B3 | 8.6±0.7 |
| | B5 | 0.98±0.09 | B5 | 8.9±0.7 |
| | B2 | 1.6±0.2 | B6 | 11±1 |
| | B6 | 1.0±0.1 | B2 | 11.6±0.9 |
| | Ch1 | 1.7±0.2 | Ch4 | 8.5±0.9 |
| | Ch4 | 2.4±0.4 | Ch5 | 10.9±0.8 |
| | Ch5 | 1.3±0.2 | Ch6 | 11.2±0.9 |
| | Ch3 | 1.7±0.3 | Ch2 | 11.7±0.8 |
| | Ch2 | 1.5±0.2 | Ch3 | 13.0±0.9 |

**Supplementary Table 5. Water-edited intensities of amino acid residues.** The intensity ratios are obtained by comparing the peak intensity in 1D $^{13}$C water-edited and control spectra, with normalization by the number of scans. Error bars are standard deviations propagated from NMR signal-to-noise ratio.

| 0.5 M NaCl | | 0 M NaCl | | 2 M NaCl | |
|---|---|---|---|---|---|
| $^{13}$C (ppm) | Intensity | $^{13}$C (ppm) | Intensity | $^{13}$C (ppm) | Intensity |
| 52.0 | 0.14± 0.09 | 52.0 | 0.2 ± 0.1 | 52.0 | 0.17 ± 0.07 |
| 43.2 | 0.12 ± 0.09 | 46.9 | 0.3 ± 0.1 | 43.2 | 0.15 ± 0.07 |
| 40.3 | 0.20 ± 0.09 | 43.2 | 0.2 ± 0.1 | 40.3 | 0.22 ± 0.07 |
| 32.6 | 0.11 ± 0.09 | 40.3 | 0.3 ± 0.1 | 32.6 | 0.14 ± 0.07 |
| 30.1 | 0.37 ± 0.09 | 30.1 | 0.5 ± 0.1 | 30.1 | 0.28 ± 0.07 |
| 27.6 | 0.27 ± 0.09 | 27.6 | 0.3 ± 0.0.1 | 27.6 | 0.16 ± 0.07 |
| 25.2 | 0.27 ± 0.09 | 25.2 | 0.4 ± 0.1 | 25.2 | 0.15 ± 0.07 |
| 19.2 | 0.16 ± 0.09 | 22.8 | 0.31 ± 0.1 | 19.2 | 0.14 ± 0.07 |
| 17.3 | 0.36± 0.09 | 19.2 | 0.2 ± 0.1 | 14.6 | 0.29 ± 0.07 |
| 14.6 | 0.34 ± 0.09 | 17.3 | 0.6 ± 0.1 | 12.0 | 0.11 ± 0.07 |
| 12.0 | 0.20± 0.09 | 14.6 | 0.6 ± 0.1 | | |
| | | 12.0 | 0.4 ± 0.1 | | |

**Supplementary Table 6. Chemical shifts of lipids.** Chemical shifts of PC and PG lipids are from measurements. The other components were from literature. Not applicable (/). Unidentified (-).

| Lipid | Carbon | $^{13}$C (ppm) | $^{1}$H (ppm) | Reference |
|---|---|---|---|---|
| PC | C2 | 34.8 | 2.3 | |
| | C3 | 25.7 | 1.6 | |
| | -CH2 | 30.5 | 1.31 | |
| | $\omega - 2$ | 32.7 | 1.3 | |
| | $\omega - 1$ | 23.4 | 1.3 | |
| | $\omega$ | 14.02 | 0.9 | |
| | α | 60.3 | 4.3 | |
| | β | 66.8 | 3.6 | |
| | γ | 54.7 | 3.2 | |
| | G1 | 63.5 | - | |
| | G2 | 71.4 | 4.4 | |
| | G3 | 63.8 | 4.0 | |
| Triglycerides | C1 | 172.4 | / | Chrissian et al. 2020[1] Lamon et al. 2023[2] |
| | C2 | 34.3 | 1.44 | |
| | C3 | 25.0 | 1.57 | |
| | $-(CH_2)_n-$ | 29.3-29.9 | 1.26-1.29 | |
| | C8,C14 | 27.3 | 1.96 | |
| | C9-C10,C12,13 | 128.1-130.0 | 5.26-5.29 | |
| | C11 | 25.6 | 2.74 | |
| | C16 | 32.0 | 1.25 | |
| | C17 | 22.8 | 1.29 | |
| | C18 | 14.1 | 0.86 | |
| | G1,G1' | 62.2 | 4.04 (α) 4.25 (β) | |
| | G2 | 69.1 | 5.18 | |
| Sterol side chain A | C20 | 36.9 | 1.33 | Suttiarporn et al. 2015[4] Chrissian et al 2020[1] |
| | C21 | 19.1 | 0.93 | |
| | C22 | 34.2 | 0.93 (α), 1.49 (β) | |
| | C23 | 30.9 | 0.93 (α), 1.37 (β) | |
| | C24 | 39.2 | 1.2 | |
| | C25 | 31.7 | 1.55 | |
| | C26 | 20.7 | 0.81 | |
| | C27 | 18.1 | 0.77 | |
| Sterol side chain B | C20 | 40.6 | 1.98 | Tuckey et al. 2012[5] |
| | C21 | 21.3 | 1.00 | |
| | C22 | 135.6 | 5.15 | |
| | C23 | 132.0 | 5.20 | |
| | C24 | 43.1 | 1.83 | |
| | C25 | 33.3 | 1.44 | |
| | C26 | 19.9 | 0.81 | |
| | C27 | 19.9 | 0.81 | |

| Lipid | Carbon | $^{13}$C (ppm) | $^{1}$H (ppm) | Reference |
|---|---|---|---|---|
| PG | C2 | 34.8 | 2.3 | |
| | C3 | 25.7 | 1.6 | |
| | -CH2 | 30.5 | 1.31 | |
| | $\omega - 2$ | 32.7 | 1.3 | |
| | $\omega - 1$ | 23.4 | 1.3 | |
| | $\omega$ | 14.02 | 0.9 | |
| | α | 63.4 | 4.1 | |
| | β | 71.1 | 4.0 | |
| | γ | 68.1 | 3.9 | |
| | G1 | - | - | |
| | G2 | 71.4 | 5.3 | |
| | G3 | - | - | |
| Polyisoprenoids | C1 trans | 39.9 | 1.94 | Chrissian et al 2020[1] |
| | C1, trans | 134.4 | / | |
| | C3 trans | 124.4 | 5.07 | |
| | C4, trans | 27.1 | 2.06 | |
| | C5, trans | 16.0 | 1.56 | |
| | C1, cis | 32.4 | 1.99 | |
| | C2, cis | 135.0 | / | |
| | C3, cis | 124.8 | 5.07 | |
| | C4, cis | 26.7 | 1.99 | |
| | C5, cis | 23.5 | 1.63 | |
| $\Delta^7$- sterol nucleus | C1 | 37.1 | 1.13(α), 1.79 (β) | Ragasa et al. 2012[3] Chrissian et al. 2020[1] |
| | C2 | 27.7 | 1.79 (α), 1.39 (β) | |
| | C3 | 73.1 | 4.62 | |
| | C4 | 34.1 | 1.69 (α), 1.28 (β) | |
| | C5 | 40.2 | 1.42 | |
| | C6 | 29.7 | - | |
| | C7 | 117.6 | 5.12 | |
| | C8 | 139.7 | / | |
| | C9 | 49.5 | 1.66 | |
| | C10 | 34.2 | / | |
| | C11 | 21.7 | 1.54 (α), 1.48(β) | |
| | C12 | 39.8 | 1.21 (α), 1.97 (β) | |
| | C13 | 43.3 | / | |
| | C14 | 55.1 | 1.80 | |
| | C15 | 23.2 | 1.55 (α), 1.97 (β) | |
| | C16 | 28.1 | 1.89 (α), 1.26 (β) | |
| | C17 | 56.3 | 1.23 | |
| | C18 | 11.9 | 0.53 | |
| | C19 | 12.9 | 0.79 | |

**Supplementary Table 7**. **Recipe of mineral-base liquid medium.** The pH is adjusted to 6.0 with $H_3PO_4$ or 0.25 M KOH. Each sample uses 100 mL of medium that contains 2 g of $^{13}C$- glucose and 0.2 g $^{15}NH_4NO_3$. The culture media and condition were adapted from a previously described protocol[6].

| Reagent | For 1 liter |
|---|---|
| $CuSO_4 \cdot 5H_2O$ | 7.8 mg |
| $FeSO_4 \cdot 7H_2O$ | 18 mg |
| $MgSO_4 \cdot 7H_2O$ | 500 mg |
| $ZnSO_4$ | 10 mg |
| KCl | 50 mg |
| $K_2HPO_4$ | 1 g |
| $^{15}NH_4NO_3$ | 2 g |
| $CuSO_4 \cdot 5H_2O$ | 7.8 mg |

**Supplementary Table 8. Solid-state NMR experiments and parameters for each of the three *A. Sydowii* samples.** To be quantitative, direct pulse (DP) experiments with 30 s long recycling delay were used. cross-polarization (CP), most rigid molecules. With DP and a shorter recycling delay of 2 seconds, suppress the rigid molecules from the spectra, and with Insensitive Nuclei Enhanced by Polarization Transfer (INEPT) the most mobile molecules were selected. For 2D $^{13}C$-$^{13}C$ correlation experiments, DARR (Dipolar Assisted Rotational Resonance) allowed to resolve rigid intramolecular peaks. While 1.5 s PDSD (Proton Driven Spin Diffusion) detects intra- and inter-molecular peaks. 2D DQ-SQ, DP J-INADEQAUTE and CP INADEQUATE spectra were used to detect through-bond correlations. The experimental parameters include the $^{1}H$ Larmor frequency, total experiment time (t), recycle delay (d1), number of scans (NS), The number of points for the direct (td2) and indirect (td1) dimensions, the acquisition time of the direct dimension (aq2) and the evolution time of indirect dimension (aq1), spectral width (sw1 and sw2), mixing time ($t_m$), increment delay (IN_F) and T filter times. * Indicates the water-polysaccharide spin diffusion and the DARR mixing time. The processing parameters include the window function and associated parameters.

| | Acquisition parameters | | | | | | | | | | | | | Processing parameters | |
|---|---|---|---|---|---|---|---|---|---|---|---|---|---|---|---|
| Experiment | $\omega_{0,\,1H}$ (MHz) | t (h) | d1 (s) | NS | td2 | td1 | aq2 (ms) | aq1 (ms) | sw2 (ppm) | sw1 (ppm) | $t_m$ (ms) | IN_F (µs) | T filters | Window function | Parameter |
| 1D CP | 850 | 0.1 | 2 | 256 | 7360 | | 14.7 | | 1169 | | | | | QSINE | SSB 3 |
| 1D DP | 850 | 0.1 | 2 | 256 | 7360 | | 14.7 | | 1169 | | | | | QSINE | SSB 3 |
| 1D DP | 850 | 2.1 | 30 | 256 | 7360 | | 14.7 | | 1169 | | | | | QSINE | SSB 3 |
| 1D INEPT | 400 | 0.1 | 3 | 64 | 3600 | | 36.0 | | 496.6 | | | | | GM | LB-5, GB0.01 |
| 1D $^{13}C$ T1 | 400 | 3.5 | 2 | 256 | 1600 | | 16.0 | | 496.6 | | | | $T_1(10^{-3}$-8 s) | | |
| 1D $^{1}H$ T1ρ | 400 | | 2 | 256 | 1400 | | 14.0 | | 496.6 | | | | SL ($10^{-3}$-30 ms) | | |
| 2D DARR | 850 | 4.2 | 2 | 48 | 1472 | 512 | 14.7 | 5.1 | 233.9 | 233.9 | 100 | 20 | | QSINE | SSB 3 |
| 2D PDSD | 850 | 13.6 | 2 | 32 | 2496 | 512 | 24.9 | 5.1 | 233.9 | 233.9 | 1500 | 20 | | QSINE | SSB 2.8 |
| 2D DP INADEQUATE | 850 | 4.5 | 2 | 32 | 7360 | 256 | 14.7 | 2.5 | 1169 | 243.4 | | 19.2 | | QSINE | SSB 3 |
| Water edited - control | 400 | 17 | 1.6 | 256 | 1400 | 152 | 14.0 | 4.9 | 496.6 | 152.8 | $10^{-4}$/50* | 65 | $T_2$ $10^{-4}$ ms | QSINE | SSB 4.5 |
| Water edited | 400 | 17 | 1.6 | 256 | 1400 | 152 | 14.0 | 4.9 | 496.6 | 152.8 | 4/50* | 65 | $T_2$ 1.2 ms | QSINE | SSB 4.5 |
| C-H INEPT | 400 | | 3.0 | 8 | 2048 | 160 | 20.0 | 11.0 | 496.6 | 18.1 | | 137 | | QSINE | SSB 3 |

**Supplementary Table 9. $^{13}C$ and $^{15}N$ chemical shifts of *A. sydowii* polysaccharides and proteins.** Superscripts are used to denote different allomorphs. Underline denotes the $^{13}C$ connectivity with ambiguity. Not applicable (/). Unidentified (-). Unknown: (Unk).

| Polysaccharides | NMR abbreviation | C1 (ppm) | C2 (ppm) | C3 (ppm) | C4 (ppm) | C5 (ppm) | C6 (ppm) | CO (ppm) | $CH_3$ (ppm) | References |
|---|---|---|---|---|---|---|---|---|---|---|
| β-1,3-glucan | B | 103.6 | 74.4 | 86.4 | 68.7 | 77.1 | 61.3 | / | / | Shim *et al.* 2007[7] |
| Chitin | Ch[a] | 103.3 | 55.5 | 72.9 | 84.5 | 76.2 | 60.7 | 174.6 | 22.7 | Fernando et al. 2021[8] |
| | Ch[b] | 103.5 | 55.2 | 73.4 | 84.4 | 75.9 | 60.0 | 173.4 | 22.7 | |
| | Ch[c] | 103.5 | 55.4 | 73.3 | 83.7 | 75.9 | 60.7 | 173.4 | 22.7 | |
| | Ch[d] | 103.6 | 55.0 | 73.2 | 83.4 | 75.5 | 60.5 | 174.1 | 22.4 | |
| | Ch[e] | 103.2 | 54,8 | 73.5 | 82.5 | 75.2 | 60.9 | 175.1 | 22.4 | |
| Chitosan | Cs[a] | 102.2 | 55.6 | 74.5 | 80.4 | 74.9 | 60.7 | / | / | |
| | Cs[b] | 101.9 | 55.7 | 72.9 | 80.0 | 74.3 | 60.5 | / | / | |
| | Cs[c] | 101.4 | 55.5 | 73.5 | 79.1 | 75.3 | 61.0 | / | / | |
| | Cs[d] | 101.4 | 55.5 | 73.5 | 79.1 | 75.3 | 61.0 | / | / | |
| α-1,3-glucan | A | 101.0 | 71.9 | 84.6 | 69.5 | 71.7 | 60.5 | / | / | Bhanja *et al.*2014[9] |
| α-1,4-galactan | Gal | 93.2 | 72.2 | 70.7 | 73.5 | 72.5 | 60.9 | / | / | Chakraborty et al 2021[10] |
| α-1,4-galactosamine | GalN | 91.7 | 54.8 | 71.1 | 81.1 | - | - | / | / | |
| α-1,4-N-acetylgalactosamine | GalNAc | 95.7 | 57.5 | 75.2 | 76.9 | - | - | - | - | |
| α-1,6-manose | Mn[1,6] | 102.7 | 70.6 | 73.2 | 72.5 | 73.7 | 66.1 | / | / | Chakraborty et al 2021[10] |
| α-1,2-manose | Mn[1,2] | 101.3 | 78.7 | 71.2 | 67.7 | 73.9 | 61.7 | / | / | |
| β-1,5 galactofuranose | Galf | 107.5 | 81.5 | 77.7 | 83.5 | 71.5 | 63.4 | / | / | |
| Unknown | Unk | 102.6 | 84.9 | 73.5 | 68.5 | - | - | | | |
| **Amino Acid** | **Abbreviation** | **Cα (ppm)** | **Cβ (ppm)** | **Cγ/γ1 (ppm)** | **Cγ2 (ppm)** | **Cδ/δ1 (ppm)** | | | | |
| Glutamic Acid | E | 55.8 | 27.8 | 34.4 | | | | | | Fritzsching. et al 2013[11] |
| Methionine | M | | 32.8 | 30.1 | | | | | | |
| Histidine | H | 55.8 | 28.2 | | | | | | | |
| Isoleucine | I | | 37.3 | 25.3 | 15.6 | | | | | |
| Arginine | R | | | 27.7 | | 40.4 | | | | |
| Cysteine | C | 55.8 | 31.0 | | | | | | | |
| Valine | V | | 30.0 | 19.2 | | | | | | |
| Leucine | L | | | 25.3 | | 22.4 | | | | |
| Alanine | A | 52.1 | 17.6 | | | | | | | |
| Proline | P | 61.7 | 30.1 | 25.8 | | | | | | |
| Lysine | K | 55.2 | 41.3 | 40.8 | 25.0 | | | | | |
